# *manta* - a clustering algorithm for weighted ecological networks

**DOI:** 10.1101/807511

**Authors:** Lisa Röttjers, Karoline Faust

## Abstract

Microbial network inference and analysis has become a successful approach to generate biological hypotheses from microbial sequencing data. Network clustering is a crucial step in this analysis. Here, we present a novel heuristic flow-based network clustering algorithm, which equals or outperforms existing algorithms on noise-free synthetic data. *manta* comes with unique strengths such as the ability to identify nodes that represent an intermediate between clusters, to exploit negative edges and to assess the robustness of cluster membership. *manta* does not require parameter tuning, is straightforward to install and run, and can easily be combined with existing microbial network inference tools.

## 1 Introduction

As most environmental covariates can only explain a small fraction of the variation in microbial communities, other factors such as species interactions have been suggested to play a large role [1]. A range of tools has become available to predict such associations. Many of these tools can predict positive as well as negative associations, with the exception of some approaches such as those based on mutual information [2]. Consequently, most microbial networks can be assigned edge weights that quantify the strength of the association. While the exact value of such edge weights may differ depending on tool usage, the sign is highly informative. Microbes can co-occur or exclude each other, as a range of ecological interactions takes place [3]. Previous work on ecological networks has demonstrated that the ratios of these interaction types can have implications for biodiversity and ecosystem stability [4–6].

While ecosystem stability, biodiversity patterns and nestedness have been linked to patterns of biotic interactions [5, 7, 8], such direct links have not been demonstrated for association networks. Unlike many ecological networks, microbial association networks suffer from interpretational challenges as they cannot be observed directly [9]. Despite these drawbacks, clusters from microbial association networks have been shown to reflect important drivers of community composition [10, 11]. However, traditional choices for network clustering algorithms are unable to make optimal use of information contained in edge signs. For example, the Markov Cluster Algorithm (MCL) uses random walks based on integers to identify network clusters [12]. While this can be achieved by scaling values or by adjusting the inflation parameter, the algorithm depends on edge density to infer clusters and is therefore mostly suitable for networks with a low number of negatively-weighted inter-cluster edges.

Alternatives have been developed that are able to take edge weight into account, such as the Louvain method for community detection [13] and the Kernighan-Lin bisection algorithm [14], an algorithm that optimizes separation of the network into two parts. An entirely different approach is to scale node weights such that negatively-weighted edges are either converted to positively-weighted edges or have lower positive weights; this approach is implemented in WGCNA [15], a pipeline for network inference and clustering. Although scaling approaches result in a loss of sign information, such approaches can also yield satisfactory results.

In this work, we describe *manta* (**m**icrobial **a**ssociation **n**e**t**work clustering **a**lgorithm), a novel flow-based method for clustering of microbial networks that can recover biologically relevant clusters. Moreover, we include a robustness metric to quantify whether cluster assignments are robust to the high error rates found in microbial association networks. Our method represents an alternative to the popular flow-based MCL algorithm [12] that in contrast to MCL can take optimal advantage of edge signs and does not need parameter optimization.

## 2 Results

### 2.1 *manta* equals or outperforms other algorithms on synthetic data sets

To evaluate performance of *manta* compared to alternative methods, we generated synthetic data sets using two different approaches. One is based on the generalized Lotka-Volterra (gLV) equation, while the other was developed for the evaluation of biclustering applied to gene expression data [16] (Fig. S2).

In the right circumstances, all algorithms achieve a separation around 0.5 when clustering the two-cluster network (Fig. 1). However, this requires the use of a positive-edge only subnetwork for the Louvain method, the WGCNA unsigned approach (data not shown) and the Girvan-Newman algorithm; on the complete network, these algorithms do not separate the nodes into two clusters or fail to separate the true-positive clusters. Shifting the edge weights failed to resolve this (Fig. S14). For MCL, separation depends on the parameter settings (Fig. S3). This algorithm can cluster the complete network, but only if the inflation parameter is set to 3 or to another uneven value. In this simulation, MCL with default parameters was unable to recover the true-positive cluster assignment (data not shown).

**Figure 1:**
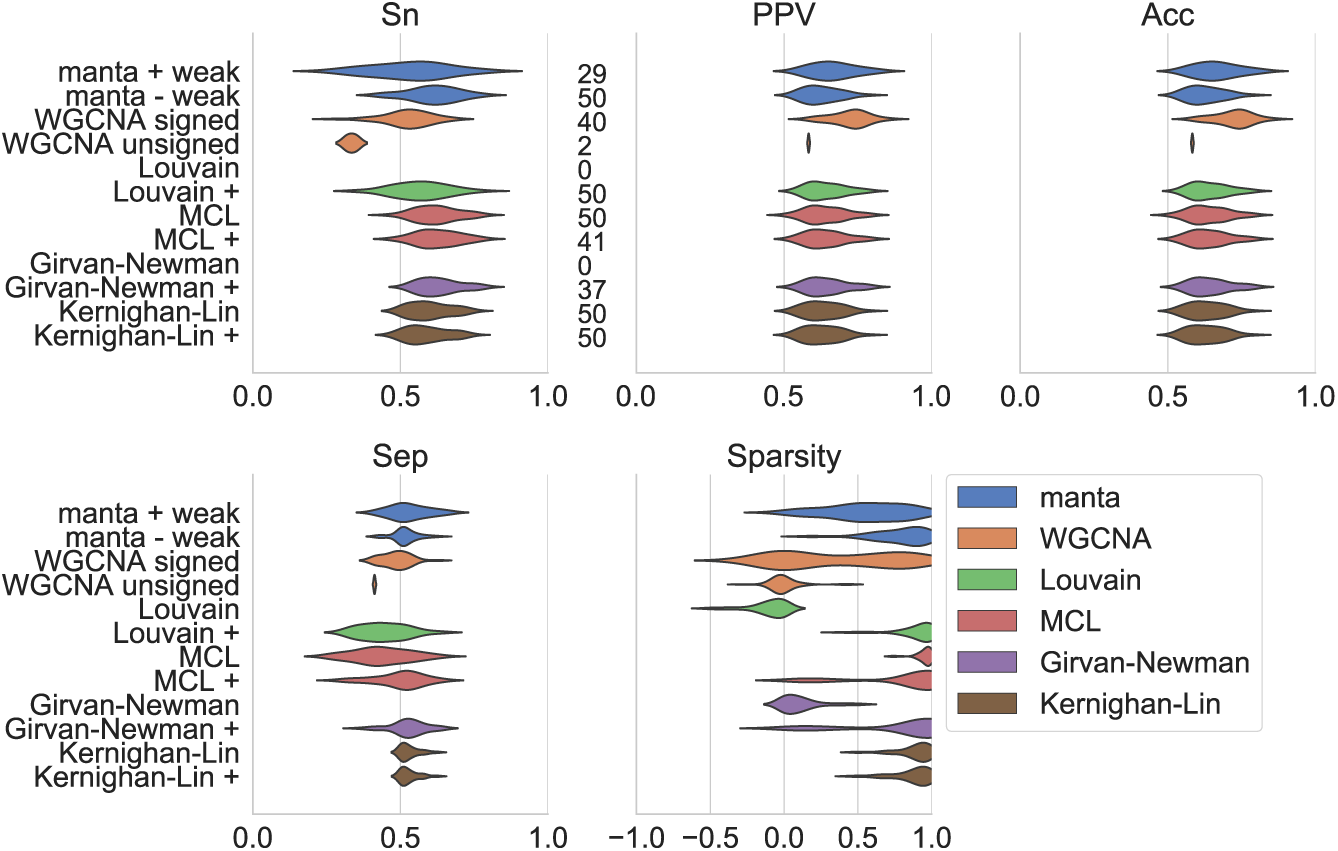
Performance of network clustering tools on two environmentally motivated clusters. Clustering performance was estimated on 50 independently generated data sets generated from random interaction matrices. Sensitivity (Sn), positive predictive values (PPV), accuracy (Acc) and separation (Sep) were calculated as described by [37]. Sparsity of the assignment is a function of the edge weights of intra-cluster versus inter-cluster edges (Equation 4). The numbers next to the sensitivity results indicate how many cluster assignments met the following criteria for a particular algorithm: no cluster should exceed 80% of the total number of nodes, and there should be fewer than 50 clusters. The *manta* algorithm was run with and without weak assignments, while WGCNA was run with signed networks and a signed topological overlap matrix and with unsigned networks combined with the unsigned matrix. For all other algorithms, we provided the complete network in addition to the positive edge-only network (indicated with +).

Strikingly, performance is improved for the Louvain method but not for the Kernighan-Lin bisection when only positively-weighted edges are considered. While the Louvain method takes edge sign into account during its optimization, negatively-weighted edges appear to have a negative effect on separation. Finally, the results for *manta* with weak assignments filtered suggests that the weak assignments increase precision even in noise-free data sets.

These environmentally motivated clusters may not necessarily reflect the true positive assignments due to species interactions amplifying or obfuscating environmental effects. Therefore, we also assessed performance on simulated biclusters generated with FABIA (Fig. 2).

**Figure 2:**
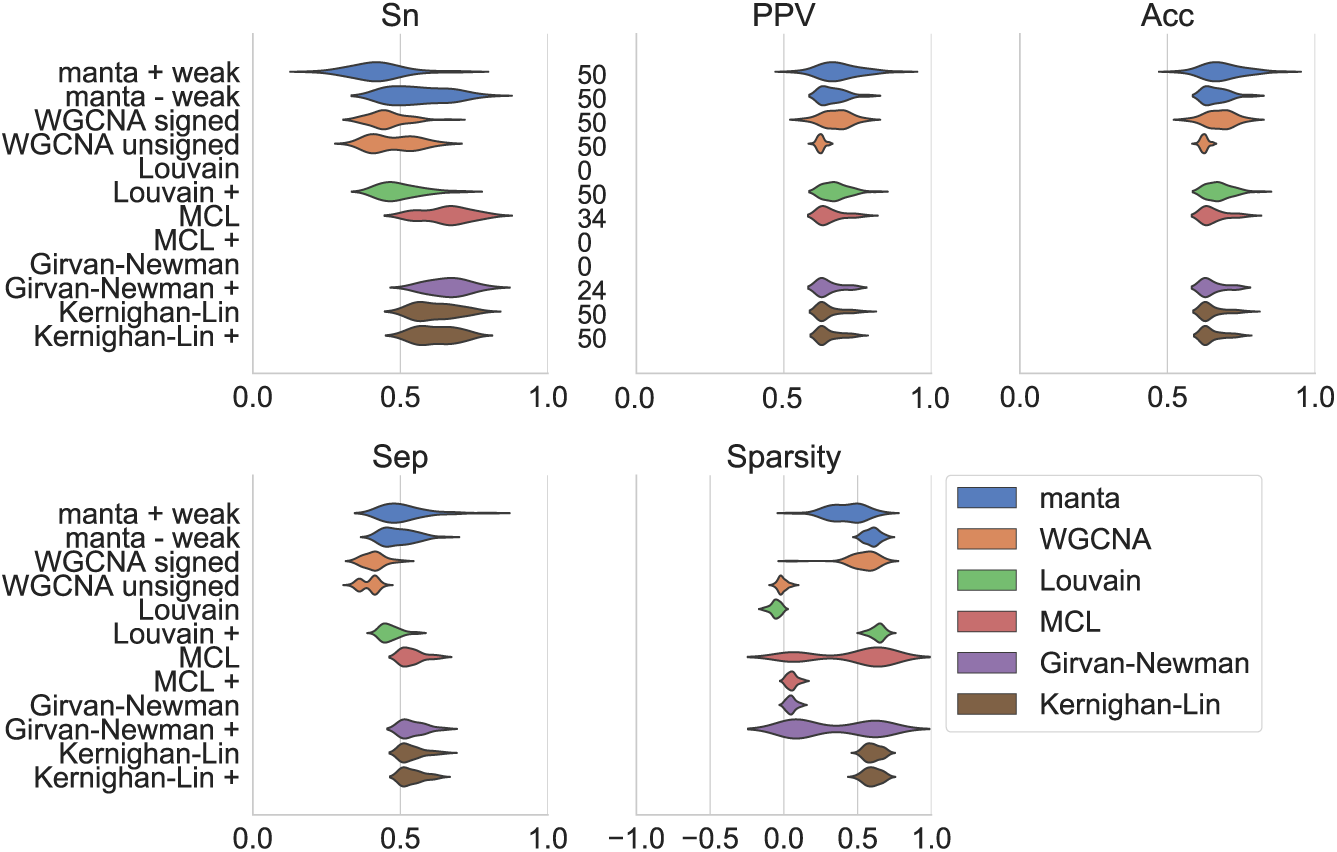
Performance of network clustering tools on two biclusters generated with FABIA [16]. Clustering performance was estimated on 50 independently generated data sets without an underlying topology. Sensitivity (Sn), positive predictive values (PPV), accuracy (Acc) and separation (Sep) were calculated as described by [37]. Sparsity of the assignment is a function of the edge weights of intra-cluster versus inter-cluster edges (Equation 4). The numbers next to the sensitivity results indicate how many cluster assignments met the following criteria for a particular algorithm: no cluster should exceed 80% of the total number of nodes, and there should be fewer than 50 clusters. The *manta* algorithm was run with and without weak assignments, while WGCNA was run with signed networks and a signed topological overlap matrix and with unsigned networks combined with the unsigned matrix. For all other algorithms, we provided the complete network in addition to the positive edge-only network (indicated with +).

The FABIA simulation has characteristics that prevent some algorithms from achieving good performance. Firstly, there is no underlying interaction network from which abundances are generated. This has a distinct effect on network topology; for example, the median approximated node connectivity of the gLV networks is 1, in contrast to 55 for the FABIA networks [17]. This implies that it is much harder to fragment the FABIA networks than the gLV networks. As a result of this change in topology, no algorithm is able to generate cluster assignments that have sparsity scores close to 1. However, not all algorithms are equally affected by the change in topology. Especially affected are those networks that assume a modular or scale-free structure, such as the Louvain method and WGCNA. The reduced performance may be attributed to one of WGCNA’s core assumptions: gene regulatory networks are assumed to be scale-free. Consequently, WGCNA infers a correlation network and soft-thresholds this network by choosing the matrix power such that scale-freeness is optimized. In the same vein, the Louvain method optimizes the modularity of cluster assignments and therefore assumes a modular structure within the data set.

The high node connectivity of the FABIA networks imply that no such modular or scale-free structure is present unless negatively-weighted edges are filtered; in that case, the Louvain method is able to return cluster assignments. Hence, approaches that imply the presence of structure are sensible for the simulation with an underlying interaction network, but may not be optimal for the FABIA biclusters. Neither *manta* nor the Kernighan-Lin method make such assumptions. As with the environmentally-motivated simulation, Kernighan-Lin bisection achieves some of the best results on this simulation. Hence, if users suspect based on a preliminary analysis that their data set contains only two clusters, this algorithm is likely to recover that separation regardless of the underlying structure. However, *manta* has the advantage of a tunable weak cluster assignment that can handle noisier data and less accurate networks and can handle data with more than 2 clusters. For MCL, parameter settings were not optimized on the FABIA data set. This dependency on parameter selection becomes apparent on data with a different structure, as the algorithm fails to recover clusters on positive-edge-only networks.

The trends described above appear to hold for 3 clusters (Supplementary Material Figs. 7, 8), for increasingly permuted data (Supplementary Material Figs. 9-12) and for networks generated from data with added multinomial noise (Fig. S13).

### 2.2 *manta* identifies biologically relevant groups in cheese rinds

We demonstrate the real-world applicability of *manta* on a cheese rind data set generated by Wolfe et al. [18]. In this study, the authors analyzed 137 cheese rind communities and identified important community members. Moreover, they found that community assembly of cheese rind communities was highly reproducible, despite the large geographical distances between cheeses. This can be explained at least partially by manipulation of the rind biofilm, as cheesemakers can introduce an initial community through starter cultures and then control the environment during the aging process. In their study on cheese rinds, Wolfe et al. [18] originally demonstrated that most of the community variation could be explained by the rind type. Indeed, most samples appear to cluster by rind type (Fig. 3B). The authors also found that samples from washed cheeses could cluster closely with both other types of rinds; the principal coordinate analysis also captures this phenomenon, as samples from washed cheeses are dispersed across the entirety of the axes.

**Figure 3:**
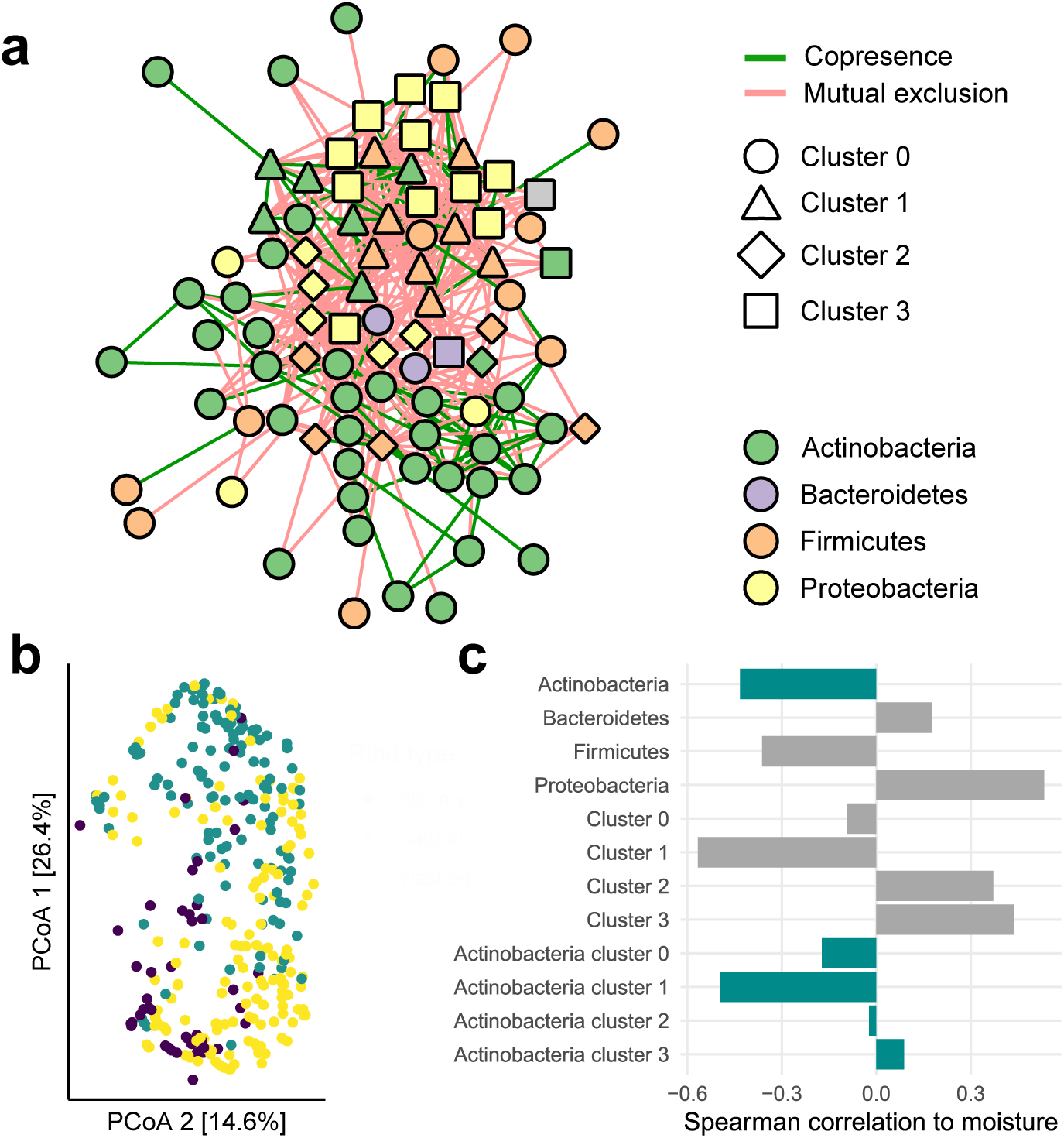
Network analysis of a cheese data set [18]. **A** CoNet network clustered with *manta*. Cluster identity is encoded in node shape, whereas node colour represents phylum membership and node border width reflects whether *manta* assigned weak cluster membership. The edge colour is mapped to the sign of the association. **B** Principal Coordinate Analysis (PCoA) of Bray-Curtis dissimilarities for sample compositions. The colours indicate the different cheese rind types: bloomy cheeses are inoculated with fungi, while washed cheeses are repeatedly washed with a brine solution. In contrast, natural rinds are not disturbed during aging. **C** Spearman correlation of moisture to summed taxon abundances. Correlations for the phylum Actinobacteria are highlighted with blue.

We ran *manta* on the association network to assess whether cluster analysis would be able to recapitulate some of the drivers of community structure in the cheeses (Fig. 3A). The network visualization of the data reveals some interesting trends, as the network contains three clusters that correlate to moisture and taxonomy. Cluster 1 is mostly comprised of Firmicutes and its summed abundances have a strong negative correlation to moisture. In fact, several of these taxa belong to the genus *Staphylococcus*, replicating the results by Wolfe et al. [18] as they demonstrated that *Staphylococcus* sp. are abundant on dry natural rinds. In contrast, cluster 1 mostly consists of Proteobacteria and correlates positively with moisture.

While the clusters correspond well to the results obtained by Wolfe et al. [18], *manta* is able to offer additional insight in community structure through its separation of Actinobacteria. Wolfe et al. [18] demonstrate that abundance of taxa belonging to this phylum is negatively correlated to moisture. However, the clusters indicate that this correlation is more nuanced. Some of the taxon abundances may reflect a gradual response to moisture rather than a strict preference for dry or moist cheese rinds, as summed abundances for Actinobacteria belonging to cluster 2 and 3 display a weak and non-significant correlation to moisture rather than the strong negative correlation associated to cluster 1.

On this data set, *manta* identifies clusters that correspond well with the main drivers of community composition in this study, while also identifying taxa that display intermediate responses to these drivers. Hence, this case study demonstrates that cluster analysis can yield novel insights into community structure.

### 2.3 Clustering global trends in coastal plankton communities

One advantage of *manta* is its ability to handle networks generated through any type of inference algorithm (though conversion to undirected networks is necessary in some cases). We demonstrate this through a time series analysis of coastal plankton communities [19]. Martin-Platero et al. [19] collected samples for 93 days and used this data to demonstrate that the communities changed rapidly, but only when lower taxonomic ranks were taken into account. The authors demonstrate WaveClust on this data set, a novel clustering method based on wavelet analysis of longitudinal data. Consequently, WaveClust can find taxon associations at low and high frequencies that are not visible without a frequency decomposition. The authors evaluated their technique on data from coastal plankton, where clusters identified by WaveClust corresponded to rapid growth of specific groups of taxa. A network analysis of the Granger causalities between clusters and metadata further demonstrated that the clusters could be separated into two regimes, both corresponding with an initial warm period followed by a rapid or gradual cooling period. These regimes are also visible in the ordination plot, as the samples separate over time around day 215 (Fig. 4B). We generated a network with eLSA and clustered this with *manta* (Fig. 4A).

**Figure 4:**
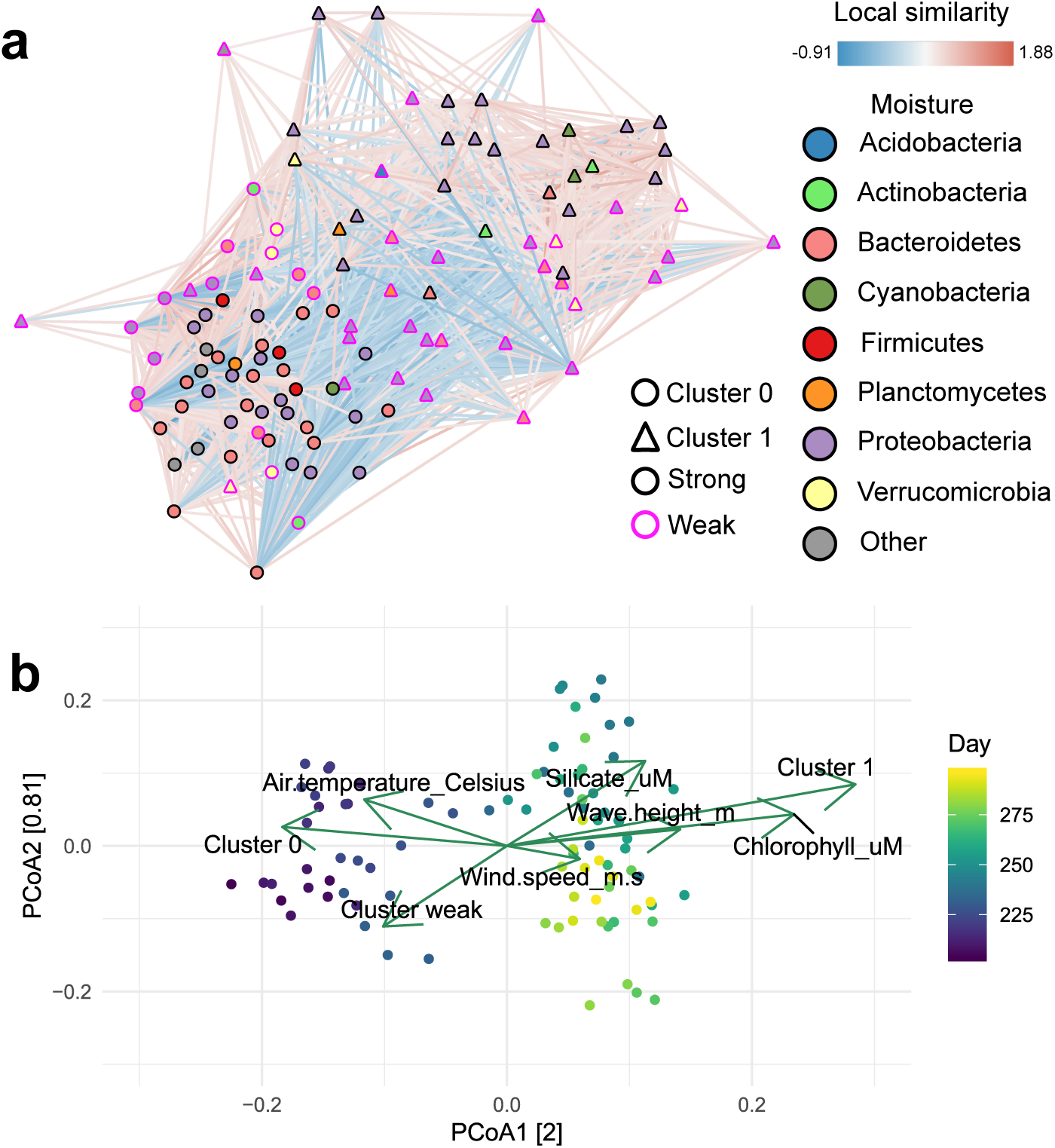
Network analysis of longitudinal 16S data collected from coastal plankton [19]. **A** eLSA network clustered with *manta*. Cluster identity is encoded in node shape, whereas node colour represents phylum membership. The edge colour is mapped to the local similarity score. **B** Principal Coordinate Analysis (PCoA) of Bray-Curtis dissimilarities for sample compositions, overlaid with environmental vectors and cluster abundance vectors. Significance of these vectors was assessed through permutation testing; only significant vectors are shown. Cluster abundance vectors were scaled independently from environmental vectors; abundances of taxa that could not be assigned to a cluster were included in the ‘Cluster weak’ vector. The axis values are the eigenvalues of the ordination axes.

*manta* is not designed for time series and therefore not able to detect frequency-specific associations. However, *manta* does summarize larger trends in the data. To demonstrate this, we first computed a network with eLSA, a microbial network inference algorithm that takes temporal shifts into account [20, 21]. A force-directed layout reveals that the eLSA network contains a large number of anti-correlated nodes. These clusters correlate with multiple environmental variables, thereby closely reflecting the two meta-regimes identified by [19]. The vector for abundances of taxa belonging to cluster 1 aligns closely with the chlorophyll concentration and wave height; in contrast, the total abundance of taxa belonging to cluster 0 was highest in the start of the experiment, when the seasonal warm period was still ongoing. The vector for taxa that could not confidently be assigned to a cluster was also significant and is directly opposite of the silicate concentration. The silicate concentration is not part of the originally reported Granger causality model [19], but it does increase sharply around the transition point between the two meta-regimes (days 240-260). Hence, the weak assignments capture a set of taxa that become less abundant during the transition period and do not correlate strongly to taxa in the assigned clusters.

## 3 Discussion

The *manta* algorithm is able to perform as well as or better than pre-existing algorithms available for network clustering, while its ability to identify weak cluster assignments can assist in defining cluster cores in networks that represent a gradient rather than distinctly separated clusters. Our case studies demonstrate how cluster assignments recapitulate main drivers of community composition.

A key limitation of *manta* is its inability to deal with networks with only a few negative edges. In such cases, cluster assignments mostly separate central nodes from peripheral nodes. However, *manta* was specifically developed for weighted networks; when only a few edges have negative weights, treating the network as unweighted or removing negatively-weighted edges will not affect network structure significantly. As demonstrated by the separation scores on positive-edge only networks, algorithms like MCL and the Louvain method will then perform adequately. Moreover, the clusters assigned by *manta* can accurately be described as ‘the enemy of my enemy is my friend’; while there are algorithms available that require nodes to be similar to nodes within the community [22], *manta* has no such requirements and may assign nodes to a cluster even though they are not necessarily positively associated to other nodes within that cluster. Additionally, users should be aware that cluster assignments will only reflect main drivers of community composition, as *manta* tends to generate a small number of clusters separated by weakly assigned nodes. Separating a data set by its main drivers (e.g. sample type or location) can help identify more interesting clusters.

Although we chose to evaluate *manta* in the context of microbial networks in this manuscript, *manta* may also be useful for clustering other types of networks with a large number of negative edges. For example, even though WGCNA was originally developed for gene expression data [15], it has also been used for microbial data [10]. The ability of *manta* to identify clusters without any underlying topological structure implies that it may be especially valuable in contexts where a small-world or scale-free structure cannot be expected. Moreover, its lack of sensitivity to parameter settings in this simulation demonstrates that *manta* is applicable in situations where little is known about the structure of the analyzed network (Supplementary Material Fig. 6).

## 4 Materials and Methods

Unless otherwise specified, computations were carried out in R v3.5.1 and Python v3.6.3. Correlation networks were generated with the *rcorr* function from the *Hmisc* R library (version 4.2-0) [23]. Analysis of simulated data sets were carried out in Python using NetworkX (version 2.1) [24], numpy (version 1.15.4) [25], pandas (version 0.21.0) [26] and scipy (version 1.2.0) [27]. Additional analyses for case studies were carried out in R using igraph (version 1.2.4.1) [28], phyloseq (version 1.26.1) [29] and vegan (version 2.5-5) [30]. In addition to the specified versions of NetworkX, numpy and scipy, *manta* uses scikit-learn (version 0.19.1) [31].

### 4.1 Clustering by graph traversal

We can represent a graph as an adjacency matrix, with each node in the graph represented as a row and column in the matrix. A non-zero entry in row *i* and column *j* represents an edge between node *i* and node *j*. In the case of an undirected graph, the adjacency matrix is symmetric. The adjacency matrix can be converted to a stochastic matrix by normalizing the values so each column sums to one. Raising the stochastic matrix *A* to a power *n* through a matrix product corresponds to the probabilities for random walks of length *n* to cross each position in the matrix. However, since this loses the sign information, we adopted a different approach that does not convert to a probability matrix.

Here, we define a scoring matrix, initialized from the weighted adjacency matrix. In this case, each position in the matrix corresponds to the edge weights. Crucially, the matrix product of the weighted adjacency matrix can take on negative values, given an even *n*. The product can then be scaled by dividing all values by the largest absolute value in the matrix (Equation 1).

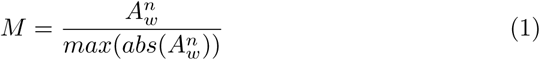

where

- *M* is the scaled scoring matrix
- *A*_*w*_ is the weighted adjacency matrix
- *n* is the walk length, by default 2

In the MCL algorithm, the separating effect of the matrix operation is boosted by raising each matrix entry to a power greater than one; afterwards, the matrix is rescaled again to acquire a column stochastic matrix (each column sums to 1) [12]. In contrast, we designed the algorithm preserve positive and negative values: every non-zero value *m* of scoring matrix *M* has its multiplicative inverse added to it (Equation 2). After inflation, the matrix is normalized again by dividing by the largest absolute value in the matrix. We also tested other variants, i.e. raising each element to a power or taking the root; unlike the multiplicative inverse, no variant was able to capture the cluster structure. While this step may seem counter-intuitive as it reverses differences rather than reinforcing them, values converge to −1 and 1 each time for the toy model (Fig. 5B).

**Figure 5:**
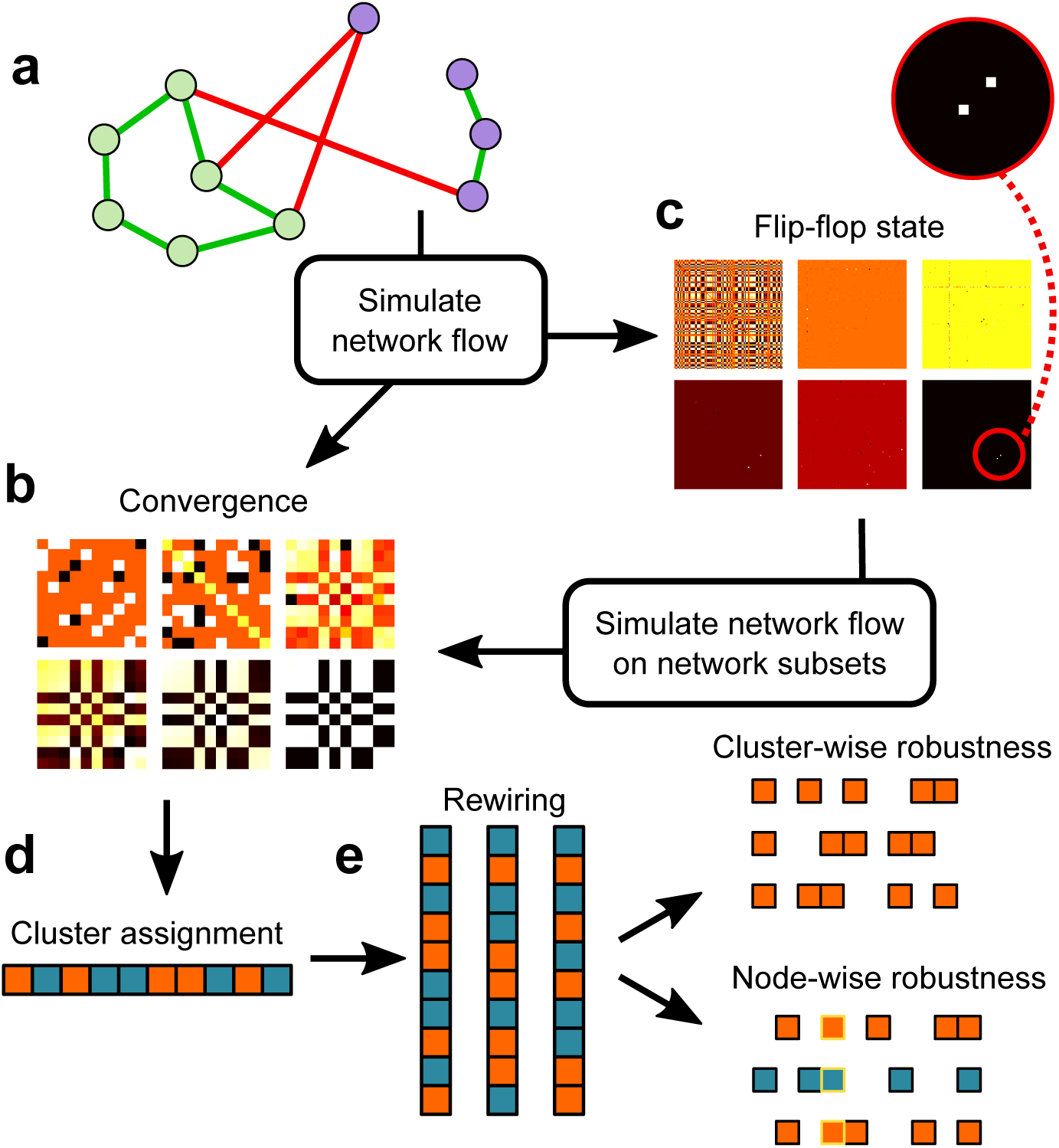
*manta* pipeline. **A** Toy graph with two clusters separated by negatively-weighted edges. **B** Scoring matrix for *A* across six iterations. Black and white values reflect −1 and 1 respectively. After six iterations, the scoring matrix reaches convergence. **C** Scoring matrix for a 100-species co-occurrence network simulated with the generalized Lotka-Volterra equation. Unlike *B*, this matrix reaches a flip-flop state. A few values in the matrix reach −1 or 1 while all other values oscillate near 0. **D** *manta* uses agglomerative clustering on the scoring matrix to assign each node to a cluster. For flip-flopping matrices, the scoring matrix is generated from subsets of the complete network. **E** A fraction of the original network is rewired to generate permuted cluster assignments with identical degree distributions. Robustness of cluster assignments can then be estimated by comparing the Jaccard similarity of cluster memberships cluster-wise or node-wise.

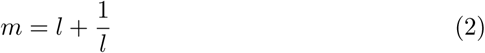

where

- *m* is an element of scoring matrix *M* after inflation before normalization
- *l* is the value of the element before inflation

The stopping condition for the algorithm is defined as a threshold ε for the average of *E* (Equation 3). The matrix of fractional differences *E* is generated from the scoring matrices of the current and the previous iterations; zero values are filtered out to prevent issues with dividing by zero.

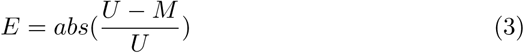

where

- *E* is the error matrix
- *U* is the updated matrix. Only positions with non-zero values are considered.
- *M* is the previous iteration of the matrix. Only positions with non-zero values in *U* are considered.

When the mean average of *E*, scaled by a factor 100 for thresholding purposes, reaches ε (e.g. 0.02), the algorithm is considered to have reached convergence. With the scoring matrix *M* as input, we use agglomerative clustering on Euclidean distances with Ward’s minimum variance method as a linkage function to define clusters [31]. The optimization criterion for choosing the optimal cluster number is based on the number of cut edges (Equation 4). For a graph *G* = (*V, e*) with edge weights *w*, the sparsity score of clusters *C*_*i*_ = (*V*_*i*_, *e*_*i*_) is calculated as a function of the sum of the cardinalities of different sets of edges based on their cluster assignment (Equation 4). Consequently, the sparsity score ranges from −1 to 1, with −1 being the worst possible cluster assignment: in this case, all negatively-weighted edges are placed inside clusters and all positively-weighted edges outside clusters.

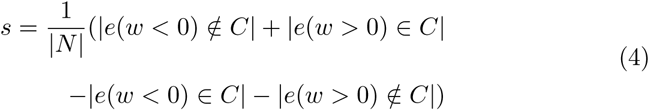

where

- *s* is the sparsity score
- *e* is an edge
- *w* is the edge weight
- *C* is the set of edges inside clusters
- *N* is the total set of edges in the graph

### 4.2 Clustering flip-flop states through a subsetting strategy

As described by Van Dongen [12], classes of matrices exist that do not converge to a stable state after repeated iterations. Instead, these matrices exhibit flip-flop equilibrium states and switch to alternative configurations with each iteration. While these flip-flop states represent rare cases when MCL is applied, the use of signed graphs by *manta* strongly increases the probability of these states appearing. For example, no convergence could be observed for larger matrices such as the simulated co-occurrence network (Fig. 5C).

This relates to the notion of balance in signed graphs [32], where a graph is only balanced if the product of edge weights in every cycle is positive. The balance of the graph matters for the expansion step; we observed that the sign of non-zero elements of the expanded scoring matrix never conflicts with the sign of non-zero elements in the weighted adjacency matrix if the adjacency matrix corresponds to a balanced graph (Supplementary Material). If the graph is not balanced [33], *manta* carries out the previously described diffusion procedure on a subset of nodes in addition to the complete network. However, only one iteration is carried out on this subset, as any more iterations would lead to the appearance of flip-flop states. Hence, this is similar to a belief propagation approach [34], where the positions in the scoring matrix are equal to the products of positions linked to adjacent edges. The propagation approach is complemented by an analysis of any balanced components in the graph; if those are present, multiple iterations of expansion and inflation are carried out on the balanced subgraph until convergence occurs. The scoring matrix used for cluster assignment is reconstructed from the subsets, but only from positions in the subsets where the sign of the value is consistent with the sign reported by most subsets. However, the accumulation of high scores for central nodes prevents the agglomerative clustering algorithm from identifying relevant clusters. Therefore, whenever small clusters with a size below a user-specified threshold are detected, rows and columns in the scoring matrix that correspond to these small clusters are removed and clusters are calculated from the remaining values. The removed nodes are then assigned to a cluster based on the average shortest path weights to cluster members.

*manta* is in theory able to handle directed graphs; in practice, strict limitations apply (Supplementary Material).

### 4.3 Weak assignments and robustness

The subsetting strategy is complemented by additional information generated through flip-flopping iterations of the network. The limits of the scoring matrix approach −1 and 1 during flip-flop iterations, but these limits are only approached by a few diagonal values. We identified a unique set of nodes corresponding to these diagonals: oscillators. The maximum of the diagonal in M approaches the positive limit for these nodes, while one position in M in the same row/column as the oscillator reaches the minimum. With the oscillators, *manta* can identify shortest paths that assess whether nodes have edges that are in conflict with their cluster assignment. For each shortest path *n* from node *v* to oscillator *t*, the product δ of the scaled edge weights is calculated. The average of these products is then the mean edge product (Equation 5).

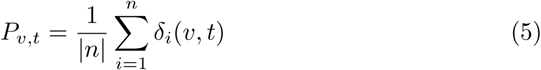

where

- *P* is the mean edge product
- *v* is a node
- *t* is an oscillator
- *n* is the set of shortest paths from *v* to *t*
- δ is the product of weights from one shortest path *i*

The node is considered to have a weak assignment if *P* meets one of the following criteria:

- The sign of *P*_*v,t*_ is not positive for *v* assigned to the same cluster as *t*.
- *P* is smaller than a user-specified threshold.

The weak assignment does not check for nodes that belong to two clusters, but rather filters out nodes that could not be assigned to any cluster, given that the shortest paths were in conflict with the node’s positions in the scoring matrix.

In addition to identification of weakly assigned nodes, we developed two new robustness measures to identify robust nodes and clusters (Fig. 5E). These measures are inspired by the reliability metric developed by Frantz and Carley [35] and highlight parts of the clustering outcome that are disproportionately sensitive to errors in network inference. Both measures are generated from rewired and are reported as a confidence interval of Jaccard similarity coefficients. For the cluster-wise robustness, Jaccard similarity scores are computed for the original clusters and their best permuted matches (identified through the maximum Jaccard similarity coefficient). This demonstrates how sensitive cluster compositions are to errors in network inference. In contrast, node-wise robustness does not use the best-matching clusters but computes Jaccard similarity coefficients for all clusters that contain the node in question.

These confidence intervals provide two different types of information. Firstly, a low Jaccard similarity coefficient indicates that a cluster has a different composition given a few errors, or that a node is assigned to a cluster with an entirely different composition given a few errors. Secondly, the width of the confidence interval demonstrates how variable cluster assignments are; wide intervals for node-wise robustness demonstrates that the node sometimes ends up in a similar cluster, but can also be assigned to a different cluster.

### 4.4 Synthetic data sets

We carried out two types of simulations with 50 replicates per simulation. For the first type, species interaction networks with a connectivity of 5% were generated with the R package seqtime (version 0.1.1). The effects of environmental factors on growth rates were sampled from a normal distribution with µ=1. The strengths of these factors were sampled from a normal distribution with µ=3. To ensure that the cumulative effects of the environmental factors were unique to each condition, one factor was set to be positively weighted while the remaining two were converted to negative values. Data sets were then generated with the generalized Lotka-Volterra equation, with growth rates of each organism adjusted per environmental condition. As some interaction matrices caused population explosions, these were re-generated until enough matrices were available for which the generalized Lotka-Volterra equation could be solved. For details and source code on these data sets, we refer to [9]; note that in contrast to this work, we did not enforce a scale-free structure in the interaction network. We generated densely connected networks from the simulated species abundances with the Pearson correlation and filtered these for significance (α=0.05). If the absolute value of the largest growth rate for one environmental condition divided by the mean growth rate was larger than 0.5, the species was assigned a specific cluster identity. This ensures that cluster assignments are not punished for failing to assign species barely affected by the environmental factors.

The second type of simulation uses an adapted version of the FABIA R package [16], a package that generates simulated data for evaluations of biclustering algorithms. We adapted the makeFabiaDatablocksPos function to generate biclusters at set locations and used these locations to define true positive clusters.

### 4.5 Clustering algorithms

We compared *manta* to the Louvain method [13], MCL [12], WGCNA [15], the Girvan-Newman method [36] and Kernigan-Lin bisection [14]; for an overview of properties relevant to this manuscript, we refer to Table 1 and the Supplementary Material. We used the following implementations of the algorithms:

**Table 1.** Overview of different clustering algorithms. Different properties of *manta*, WGCNA, MCL, Louvain community detection, the Girvan-Newman algorithm and the Kernighan-Lin bisection algorithm. The following properties are summarized: how algorithms choose a cluster number, whether they can leave nodes unassigned, whether they perform better with negatively-weighted edges removed and what types of networks they accept. Finally, we assessed whether algorithms required extensive parameter tuning before achieving optimal performance on simulated data.

| Algorithm | Cluster number criterion | Unassigned nodes | No edge filtering required | Preferred network type | No parameter tuning |
| --- | --- | --- | --- | --- | --- |
| *manta* | Optimize sparsity | ✓ | ✓ | Undirected | ✓ |
| WGCNA signed | Dynamic branch cut | ✓ | ✓ | None (constructs network) | ✓ |
| WGCNA unsigned | Dynamic branch cut | ✓ | X | None (constructs network) | ✓ |
| Louvain method | Optimize modularity | X | ✓ | Undirected; directed version possible | X |
| MCL | Parameter-dependent | X | ✓ if parameters optimized | Undirected | X |
| Girvan-Newman algorithm | User-dependent | X | X | Undirected; directed version possible | ✓ |
| Kernighan-Lin bisection | Only bisection | X | ✓ | Any | ✓ |

- WGCNA: blockwiseModules function (version 1.66) [15]
- MCL: markov_clustering (version 0.0.5) (https://github.com/GuyAllard/markov_clustering)
- Louvain method: python-louvain (version 0.11) (https://github.com/taynaud/python-louvain)
- Girvan-Newman algorithm: networkx (version 2.1)
- Kernighan-Lin bisection: networkx (version 2.1)

We supplied both the complete network and the positive-edge only network to each algorithm except WGCNA and *manta*. WGCNA received the simulated data set instead of the Pearson correlations. For its evaluation, only assigned nodes were considered (i.e. nodes with poor correlations to cluster eigenvectors were ignored). A range of parameter settings was tested for *manta*, the Louvain method, the Kernighan-Lin algorithm and MCL (Supplementary Material Figs. 3–6). We set these parameters as follows:

- *manta*: ratio set to 0.8, edgescale to 0.3
- MCL: On complete networks, inflation was set to 3 and expansion to 15. On positive-edge-only networks, inflation was set to 2 and expansion to 7.
- Louvain method: On complete networks, resolution was set to 0.1. On positive-edge-only networks, resolution was set to 1.
- Girvan-Newman algorithm: no parameter settings.
- Kernighan-Lin bisection: On all networks, max_iter was set to 10.

Cluster assignments were evaluated as described by [37]. We report both the complex-wise sensitivity (Sn), the cluster-wise positive predictive value (PPV), geometrical accuracy (Acc) and the separation (Sep) (Fig. S1). We also report the sparsity score (Equation 4). When a cluster size exceeded 80% of the total number of assigned species, all measures except the sparsity score were replaced with missing values to avoid obfuscation of the results. The opposite case - when nearly all species were assigned to their own clusters - was also replaced when the number of assigned clusters exceeded 50.

### 4.6 Cheese rind case study

We downloaded BIOM-formatted bacterial abundances of this cheese data set (study ID 11488, [18]) from Qiita [38]. The data set was preprocessed by removing samples with fewer than 10000 counts, rarefying to even depth and filtering taxa with less than 20% prevalence. The final data set therefore included 97 taxa and 337 samples. We used these abundances to construct a network with CoNet; an initial multigraph was constructed with Pearson correlation, Spearman correlation, mutual information, Bray-Curtis dissimilarity and Kullback-Leibler dissimilarity. Only edges that were supported by at least 2 methods were retained. The bootstrapping procedure further removed edges with p-values below 0.05, where p-values were merged with Brown’s method and the Benjamini-Hochberg correction for multiple testing was applied. For clustering with *manta*, edge weights were converted to −1 or 1 based on the inferred sign. Additionally, we inferred Spearman correlations of taxon abundances to moisture, where taxon abundances were either summed per phylum or per cluster. *manta* was run with default settings (edge scale and ratio set to 0.8).

### 4.7 Longitudinal coastal plankton case study

We downloaded the supplementary files from [19]. Taxon abundances were first agglomerated at genus level; afterwards, we removed samples with fewer than 2000 counts and rarefied to even depth. Taxa with a sample prevalence below 30% were removed, with the total counts preserved by including them in a bin. This resulted in a final dataset of 143 taxa and 260 samples. Removed samples were given as columns with NA values, and the time series supplied to eLSA therefore contained 90 timepoints with 3 replicates per timepoint. We then ran eLSA (v1.0.2, Python v2.7.12) with simple replicate merging (averaging replicates) and a delay of 5 [20, 21]. For downstream analysis with *manta*, only associations with p-values below 0.05 and q-values below 0.05 were taken into account. The network was treated as undirected and edge weights were converted to −1 and 1 prior to clustering. *manta* was run with default settings except for an edge scale of 0.2; with the default edge scale, nearly all nodes in the network were determined to be weakly assigned.

For an additional analysis of the clusters, a principal coordinate analysis of the metadata with Bray-Curtis dissimilarity was carried out. Technical replicates were averaged and the metadata features supplied by [19] were used to fit environmental vectors onto the ordination. Significance of these vectors was assessed through permutation testing (1000 permutations), and only vectors with a p-value below 0.05 (after Benjamini-Hochberg multiple testing correction) and q-value below 0.05 were retained. To compare the direction of cluster abundances to these vectors, all bacterial abundances were summed and the covariance between these abundance vectors and principal component vectors estimated. These covariance values significantly exceeded covariances of permuted bacterial abundances (1000 permutations, p-value 0.001) and were used to construct cluster abundance vectors; an arbitrary scaling coefficient was used for visualization purposes.

## 5 Data Availability

All code for *manta* is available under the Apache-2.0 license at https://github.com/ramellose/manta. An archived version of *manta*, together with code and synthetic data used for writing the manuscript, is available from https://zenodo.org/record/3407044#.XXtTcmZx1EY.

## 6 Acknowledgements

We would like to acknowledge Didier Gonze for helpful feedback on the balance of signed matrices. This work was financially supported by KU Leuven.

