## Supplementary Materials for "*manta* - a clustering algorithm for weighted ecological networks"

### Contents

**1 Balanced graphs and sign changes in the scoring matrix** **3**

**2 Directed graphs** **3**

**3 Evaluation of clustering performance** **3**

**4 Effect of network errors on algorithm performance** **4**

**5 Supplementary Figures** **5**

### 1 Balanced graphs and sign changes in the scoring matrix

A balanced graph can only contain cycles where the product of all edge weights is positive. Repeating the cycle twice or more does not change the sign, while a repetition of an unbalanced cycle does change the sign of the path for each repetition. While cycles are only a subset of the paths that are included in the matrix power, the ratio of paths that switch signs is different for balanced and unbalanced graphs as a result of their presence. Hence, we hypothesize that this is what causes the oscillating behaviour in unbalanced, but not in balanced graphs.

### 2 Directed graphs

During iterations on directed graphs, the scoring matrix can also be described as a Petri net, where positions in the scoring matrix can be assigned a liveness value based on their ability to affect the matrix throughout iterations. While *manta* does support directed graphs, it can only cluster these when the liveness of some matrix positions is at least $L_2$, where the $L_2$ liveness of positions implies that they generate values other than zero through multiple iterations of expansion and inflation [1]. The occasional non-zero values generated for permuted graphs with $L_2$ liveness could be enough to assign clusters with the subsetting strategy; however, random selection of nodes does not enforce $L_2$ liveness. Therefore, the scoring matrix needs to converge to -1 and 1 during the initial cluster assignment and no convergence to zero is permitted.

As the subsetting strategy does not enforce a degree of liveness on subsets of directed graphs, *manta* cannot resolve cluster structure for flip-flopping directed graphs. Moreover, most of the use cases for *manta* are unlikely to enforce a degree of liveness on the input network. Hence, we recommend users to treat directed networks as undirected instead.

### 3 Evaluation of clustering performance

Cluster assignments were evaluated with the complex-wise sensitivity (Sn), the cluster-wise positive predictive value (PPV), geometrical accuracy (Acc) and the separation (Sep) [2]. The complex-wise sensitivity estimates the coverage of a true positive cluster by its best-matching assigned cluster, whereas the cluster-wise positive predictive value measures how well an assigned cluster covers its best-matching true positive cluster. In contrast, the separation is calculated by taking the product of the fraction of assigned nodes in the true positive clusters by the fraction of true-positive nodes in the assigned clusters. Hence, the separation penalizes for cluster overlap, unlike the reported Acc, PPV and Sn. The approach described in uses the contingency matrix rather than a list of true positives, effectively permitting evaluation of assignments that do not necessarily match the true positive clusters (Supplementary Figure 1). These measures can be skewed by cluster assignments that mostly assign all nodes to one cluster (Supplementary Figure 1b) or assign almost every node to its own cluster. Additionally, overlapping clusters can also inflate some measure of performance (Supplementary Figure 1c). In the first and second cases, all or some measures are inflated; therefore, we filtered assignments where over 80% of the nodes were assigned to a single cluster, or over 50 clusters were identified. While accuracy, precision and sensitivity can be high for algorithms that assign true positive clusters to the same clusters, separation is calculated by multiplying the proportion of true-positive nodes in the assigned cluster with the proportion of cluster nodes in the true-positive cluster. Hence, the separation measure punishes cluster assignments that mix up multiple true-positive clusters. Finally, the reported sparsity is a measure of the ratio between inter- and intracluster positive and negative edges.

### 4 Effect of network errors on algorithm performance

One of the main conclusions of the work by Brohee et al. [2] was that MCL was exceptionally robust to alterations of the network. In a similar manner, we carried out an evaluation to test algorithms for robustness to alterations of the abundance data. We permuted the original abundances generated with the gLV and FABIA approaches to assess whether algorithms were able to capture the original clusters (Supplementary Figure 9). For each algorithm, the best-performing variant on the population model was chosen (Figure 2), i.e. the positive-edge-only assignment for the Louvain method and signed approach for WGCNA. Except WGCNA, all algorithms show a decrease in separation after a large fraction of the data is permuted. Although all algorithms, including manta, display a decrease in performance, they still recover at least part of the original clusters despite a high degree of errors. Considering the fraction of permuted values, it is likely that they are mostly identifying a random cluster structure. Regardless, the Kernighan-Lin algorithm and *manta* both recover part of the true positive clusters, with *manta* outperforming all other algorithms on data sets with 3 clusters (Supplementary Figures 10, 12). On data with multinomial noise, the performance of most algorithms was hardly affected (Supplementary Figure 13).

#### 5 Supplementary Figures

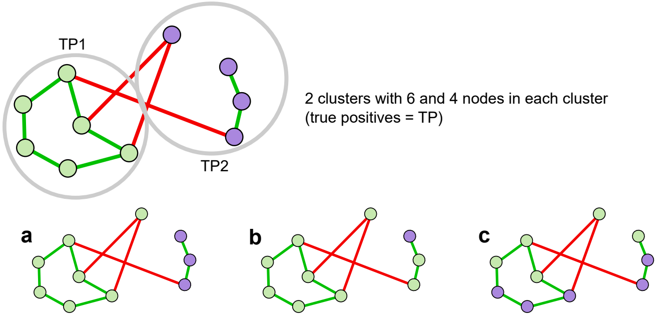


**Contingency matrix**

| | a A1 | a A2 | b A1 | b A2 | c A1 | c A2 |
|---|---|---|---|---|---|---|
| TP1 | 6 | 0 | 6 | 0 | 3 | 3 |
| TP2 | 1 | 3 | 3 | 1 | 2 | 2 |

**Sensitivity (complex-wise) value / row sum**

| | a A1 | a A2 | b A1 | b A2 | c A1 | c A2 |
|---|---|---|---|---|---|---|
| TP1 | **1** | 0 | **1** | 0 | 0.5 | 0.5 |
| TP2 | 0.25 | **0.75** | **0.75** | 0.25 | 0.5 | 0.5 |

| | a | b | c |
|---|---|---|---|
| Sensitivity (**cluster-wise**) weighted average | **0.9** | **0.9** | **0.5** |

**Positive predictive value (complex-wise) value / column sum**

| | a A1 | a A2 | b A1 | b A2 | c A1 | c A2 |
|---|---|---|---|---|---|---|
| TP1 | **0.86** | 0 | **0.66** | 0 | **0.6** | **0.6** |
| TP2 | 0.14 | **1** | 0.33 | **1** | 0.4 | 0.4 |

| | a | b | c |
|---|---|---|---|
| Positive predictive value (**cluster-wise**) weighted average | **0.9** | **0.7** | **0.6** |
| Accuracy Geometrical mean | **0.9** | **0.79** | **0.55** |

**Separation Row frequency * Column frequency**

| | a A1 | a A2 | b A1 | b A2 | c A1 | c A2 |
|---|---|---|---|---|---|---|
| TP1 | 0.86 | 0 | 0.66 | 0 | 0.3 | 0.3 |
| TP2 | 0.04 | 0.75 | 0.25 | 0.25 | 0.2 | 0.2 |

| | a | b | c |
|---|---|---|---|
| Separation Geometrical mean of cluster-wise and column-wise separation | **0.82** | **0.58** | **0.5** |

Figure S1: **Effect of different clustering assignments on measures for clustering performance.** Overview of sensitivity, positive predictive values, accuracy and separation as described by [2]. A toy model contains two true positive clusters (TP1 and TP2); the effect of clustering assignments (A1 and A2) on these measures is shown in tables. **a** A clustering assignment that assigns 9 out of 10 nodes correctly achieves good scores for all measures. **b** A clustering assignment that assignes 9 out of 10 nodes to the same cluster achieves high sensitivity, but lower values for all other scores. **c** A clustering assignment that incorrectly assigns most nodes achieves values of approximately 0.5-0.6 for all measures.

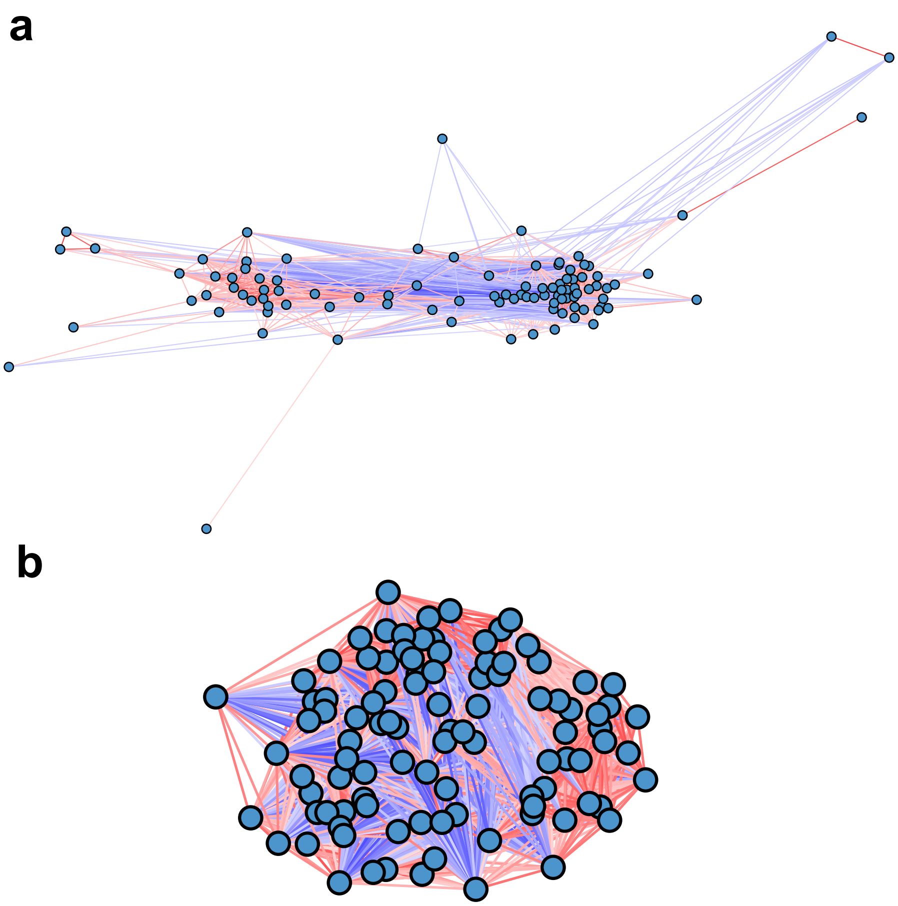


Figure S2: **Examples of networks generated with generalized Lotka-Volterra or FABIA.** The edge colours are mapped to the Spearman correlation, with red being a positive and blue a negative correlation. **a** A network generated with the generalized Lotka-Volterra equation from a random interaction matrix. **b** A network generated with the FABIA package [3].

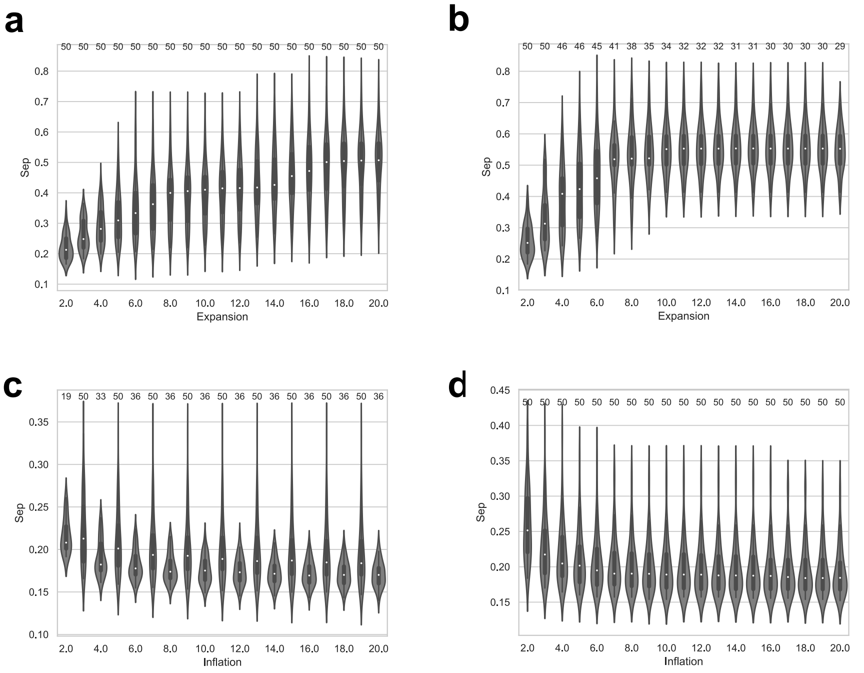


Figure S3: **Effect of MCL parameters on clustering performance.** Separation of MCL clustering assignments. Parameter choices for the MCL algorithm were based on performance on datasets generated with random interaction matrices and two environmentally-induced clusters. Cluster assignments that assigned over 80% of species to one cluster are not shown. The numbers above each setting indicate how many cluster assignments without clusters larger than 80% of the dataset were returned. **a** Effect of the expansion variable on separation of complete networks. The inflation parameter was set to 3. **b** Effect of expansion variable on separation of positive-edge-only networks. The inflation parameter was set to 2. **c** Effect of the inflation variable on separation of complete networks. The expansion variable was set to 2. **d** Effect of the inflation variable on separation of positive-edge-only networks. The expansion variable was set to 2.

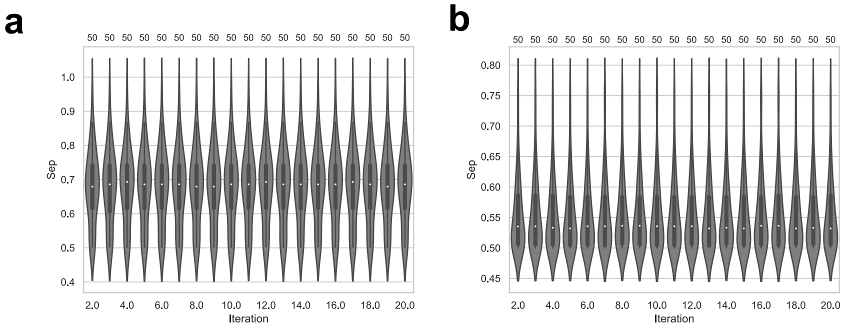


Figure S4: **Effect of Kernighan-Lin parameters on clustering performance.** Separation of Kernighan-Lin clustering assignments. Parameter choices for the Kernighan-Lin algorithm were based on performance on datasets generated with random interaction matrices and two environmentally-induced clusters. The numbers above each setting indicate how many cluster assignments without clusters larger than 80% of the dataset were returned. **a** Effect of the iteration variable on separation of complete networks. **b** Effect of the iteration variable on separation of positive-edge-only networks.

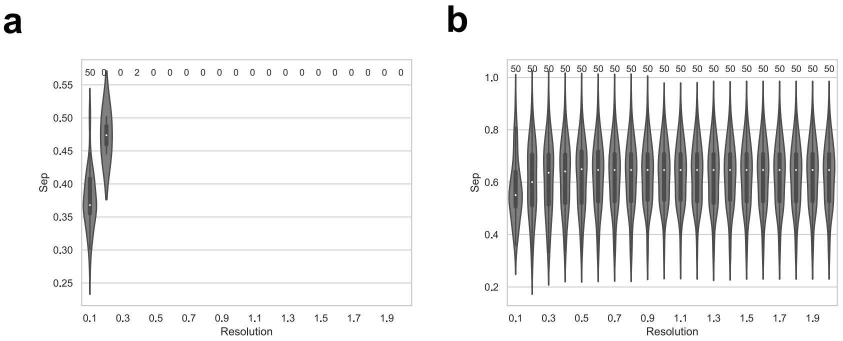


Figure S5: **Effect of Louvain community detection parameters on clustering performance.** Separation of Louvain community assignments. Parameter choices for the Louvain method for community detection were based on performance on datasets generated with random interaction matrices and two environmentally-induced clusters. Cluster assignments that assigned over 80% of species to one cluster are not shown. The numbers above each setting indicate how many cluster assignments without clusters larger than 80% of the dataset were returned. **a** Effect of the resolution variable on separation of complete networks. **b** Effect of the resolution variable on separation of positive-edge-only networks.

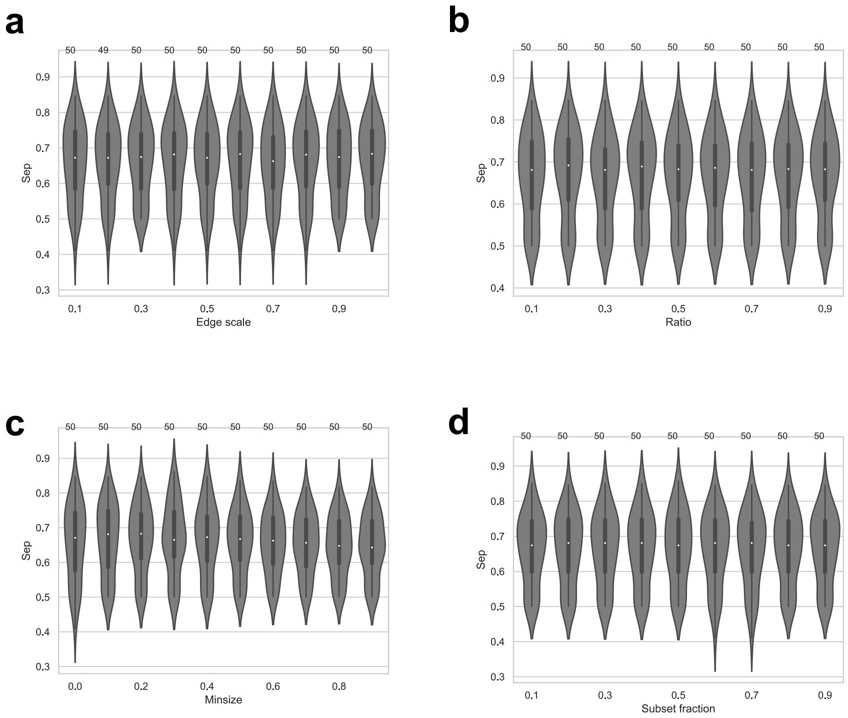


Figure S6: **Effect of *manta* clustering parameters on clustering performance.** Separation of *manta* clustering assignments. Parameter choices for the *manta* algorithm were based on performance on datasets generated with random interaction matrices and two environmentally-induced clusters. The numbers above each setting indicate how many cluster assignments without clusters larger than 80% of the dataset were returned. **a** Effect of the edgescale variable on complete networks. **b** Effect of the ratio variable on complete networks. **c** Effect of the minsize variable on separation of complete networks. **d** Effect of the subset variable on separation of complete networks.

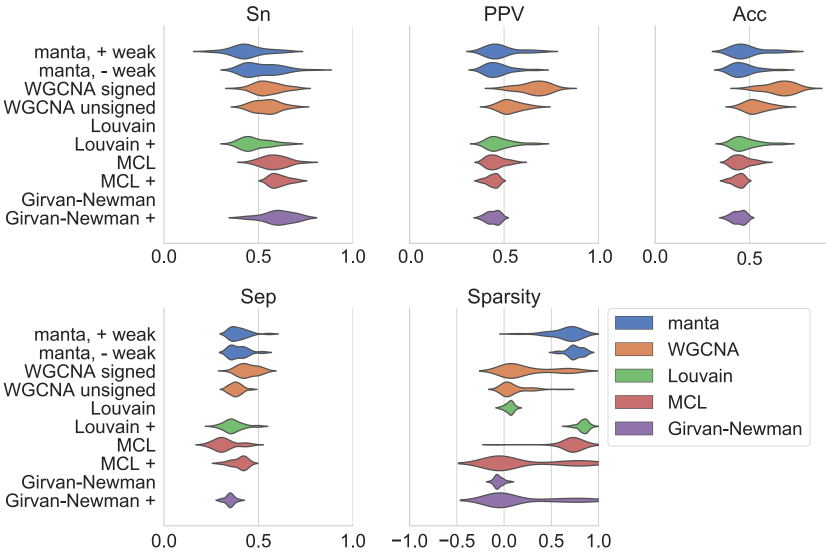


Figure S7: **Performance of network clustering tools on three environmentally motivated clusters.** Clustering performance was estimated on 50 independently generated datasets generated from scale-free interaction matrices. Sensitivity (Sn), positive predictive values (PPV), accuracy (Acc) and separation (Sep) were calculated as described by [2]. Sparsity of the assignment is a function of the edge weights of intra-cluster versus inter-cluster edges (Equation 4). The *manta* algorithm was run with and without weak assignments, while WGCNA was run with signed networks and a signed topological overlap matrix and with unsigned networks combined with the unsigned matrix. For all other algorithms, we provided the complete network in addition to the positive edge-only network (indicated with +).

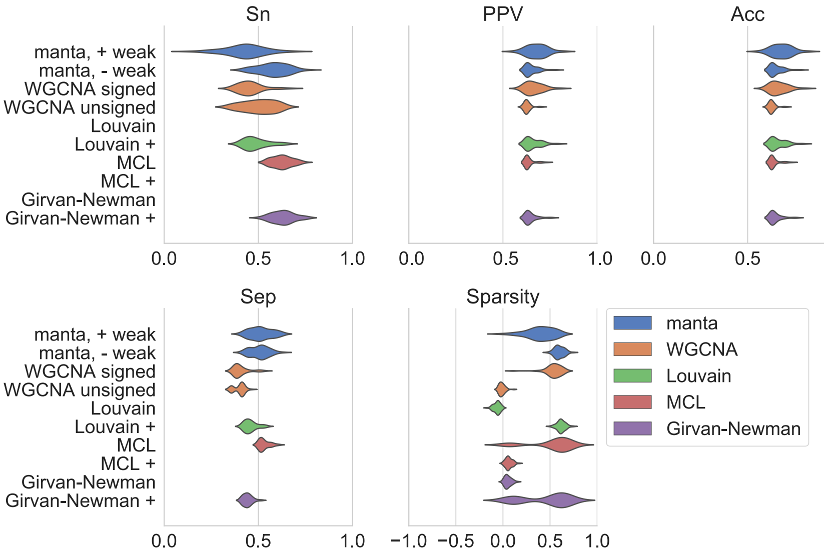


Figure S8: **Performance of network clustering tools on three biclusters generated with FABIA [3].** Clustering performance was estimated on 50 independently generated datasets without an underlying topology. Sensitivity (Sn), positive predictive values (PPV), accuracy (Acc) and separation (Sep) were calculated as described by [2]. Sparsity of the assignment is a function of the edge weights of intra-cluster versus inter-cluster edges (Equation 4). The *manta* algorithm was run with and without weak assignments, while WGCNA was run with signed networks and a signed topological overlap matrix and with unsigned networks combined with the unsigned matrix. For all other algorithms, we provided the complete network in addition to the positive edge-only network (indicated with +).

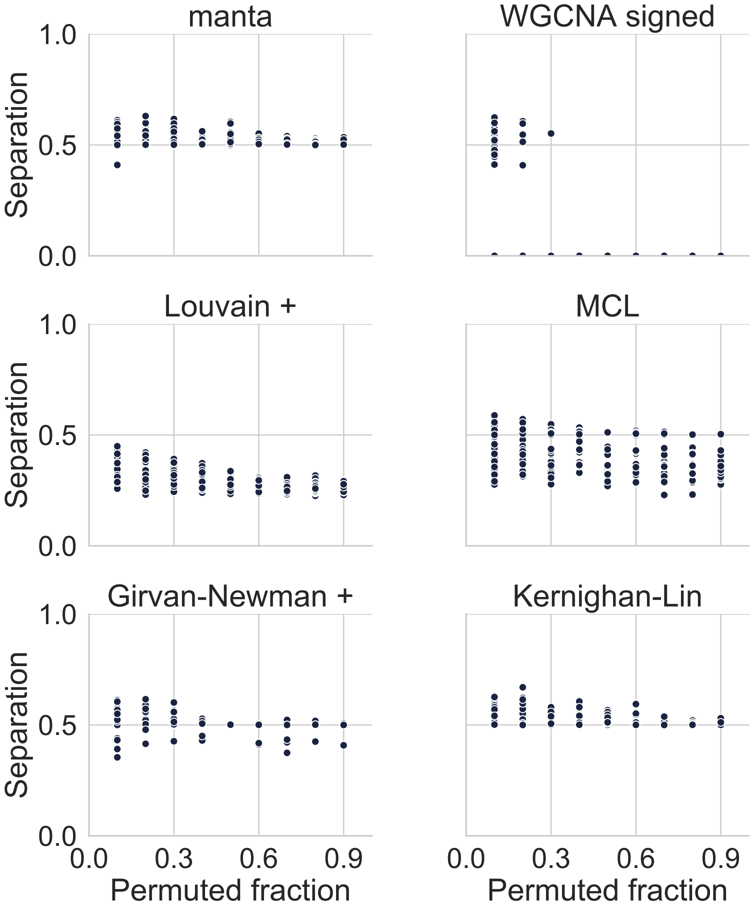


Figure S9: **Performance of clustering algorithms across a permutation gradient.** Clustering performance was estimated on 50 independently generated data sets using the separation as described by [2]. For 9 points (range 0.1-0.9), fractions of the original abundance matrices (generated from a random interaction matrix with two environmentally-induced clusters) were permuted and clustering was carried out on Pearson correlation networks inferred from these matrices. The *manta* algorithm was run without weak assignments, while WGCNA was run with signed networks and a signed topological overlap matrix. The Louvain method and Girvan-Newman algorithm received positive-edge-only networks (indicated with +), while MCL received the complete network.

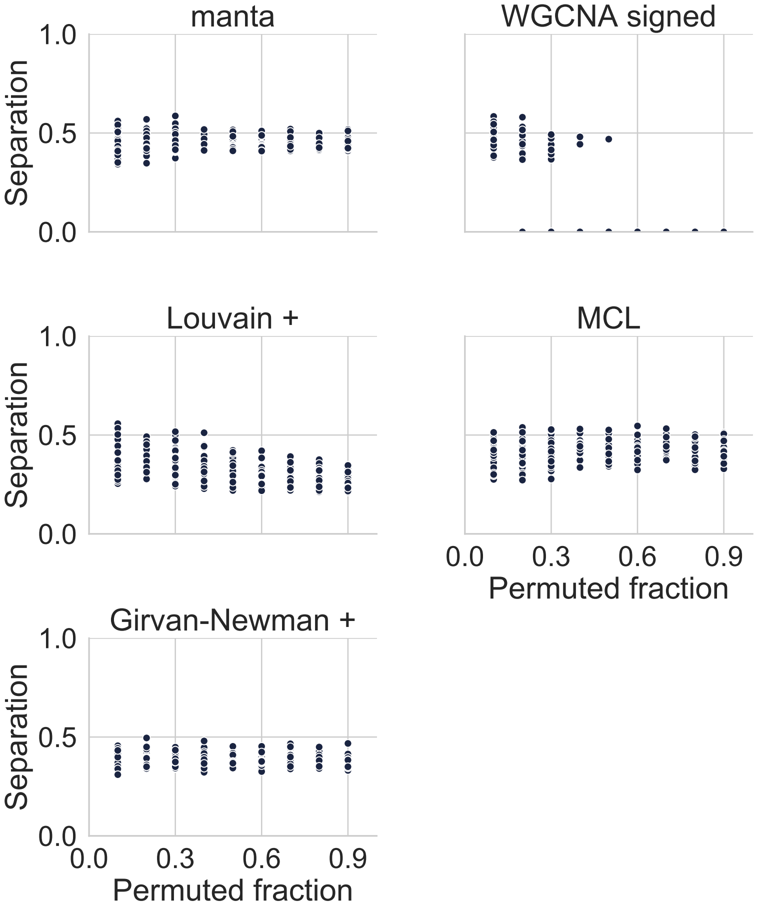


Figure S10: **Performance of clustering algorithms across a permutation gradient on three environmentally motivated clusters.** Clustering performance was estimated on 50 independently generated datasets using the separation as described by [2]. For 9 points (range 0.1-0.9), fractions of the original abundance matrices (generated from a random interaction matrix with two environmentally-induced clusters) was permuted and clustering was carried out on Spearman correlation networks inferred from these matrices. The *manta* algorithm was run without weak assignments, while WGCNA was run with signed networks and a signed topological overlap matrix. The Louvain method and Girvan-Newman algorithm received positive-edge-only networks (indicated with +), while MCL received the complete network.

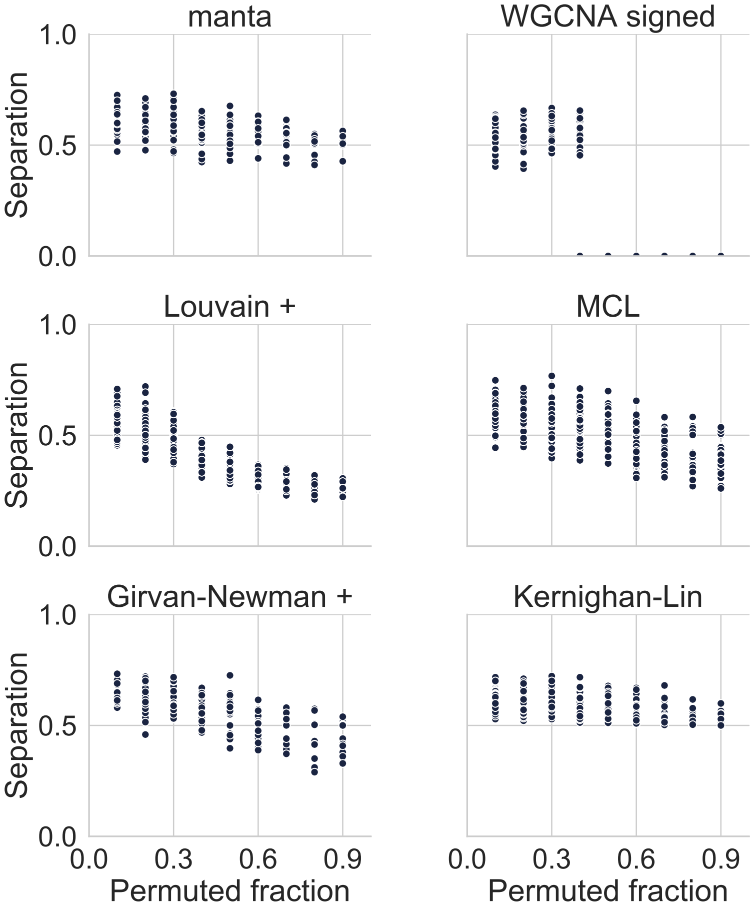


Figure S11: **Performance of clustering algorithms across a permutation gradient on two biclusters generated with FABIA [3].** Clustering performance was estimated on 50 independently generated datasets using the separation as described by [2]. For 9 points (range 0.1-0.9), fractions of the original abundance matrices (generated from a random interaction matrix with two environmentally-induced clusters) was permuted and clustering was carried out on Spearman correlation networks inferred from these matrices. The *manta* algorithm was run without weak assignments, while WGCNA was run with signed networks and a signed topological overlap matrix. The Louvain method and Girvan-Newman algorithm received positive-edge-only networks (indicated with +), while MCL received the complete network.

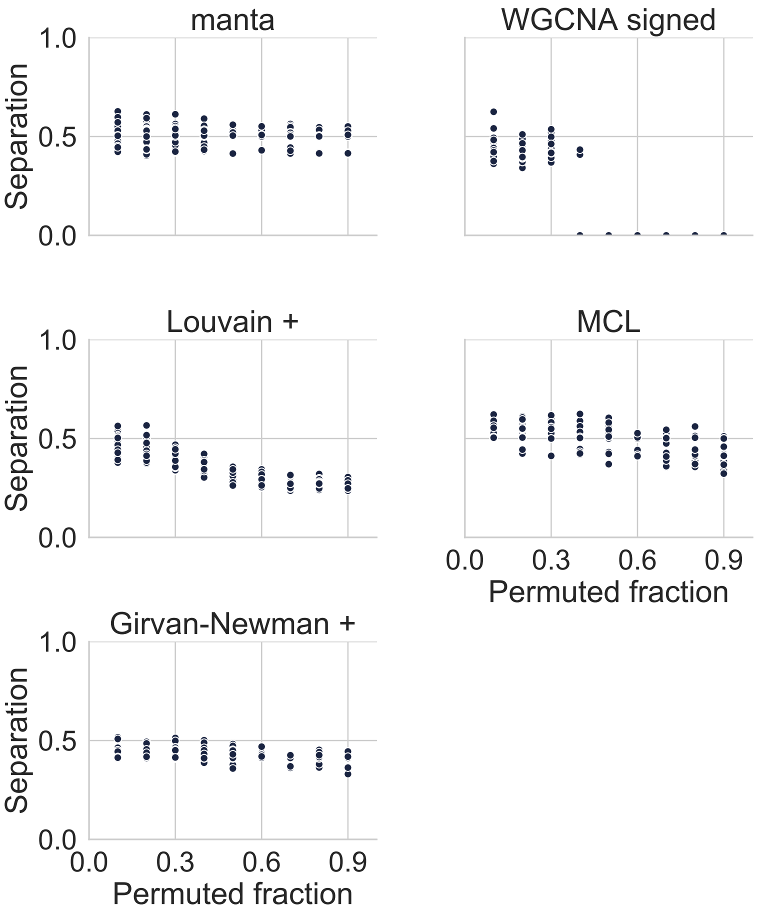


Figure S12: **Performance of clustering algorithms across a permutation gradient on three biclusters generated with FABIA [3].** Clustering performance was estimated on 50 independently generated datasets using the separation as described by [2]. For 9 points (range 0.1-0.9), fractions of the original abundance matrices (generated from a random interaction matrix with two environmentally-induced clusters) was permuted and clustering was carried out on Spearman correlation networks inferred from these matrices. The *manta* algorithm was run without weak assignments, while WGCNA was run with signed networks and a signed topological overlap matrix. The Louvain method and Girvan-Newman algorithm received positive-edge-only networks (indicated with +), while MCL received the complete network.

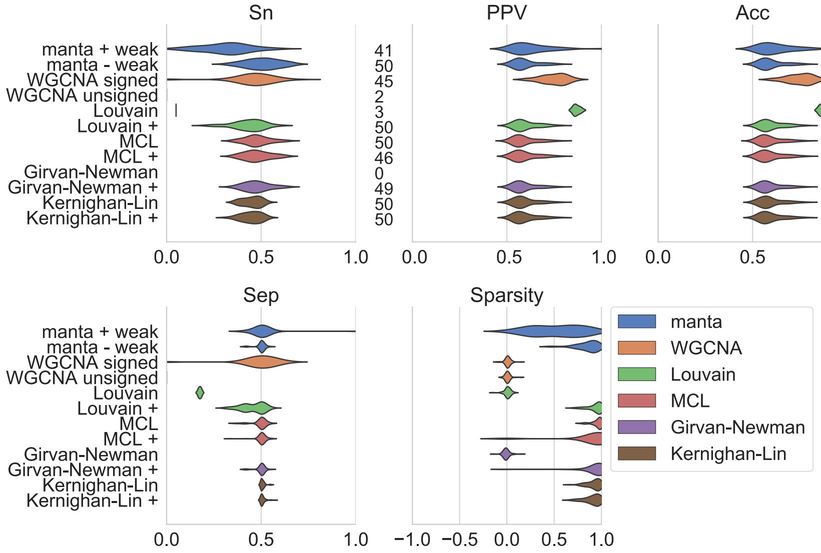


Figure S13: **Performance of clustering algorithms on networks generated from noisy data.** Sensitivity (Sn), positive predictive values (PPV), accuracy (Acc) and separation (Sep) were calculated as described by [2]. Sparsity of the assignment is a function of the edge weights of intra-cluster versus inter-cluster edges (Equation 4). Clustering performance was estimated on 50 independently generated datasets. Matrices of taxon abundances were generated from a synthetic random interaction matrix; afterwards, taxon abundances were scaled by a factor 1000 and multinomial noise was applied. Clustering was carried out on Spearman correlation networks inferred from these matrices. The numbers next to the sensitivity results indicate how many clustering assignments met the following criteria for a particular algorithm: no cluster should exceed 80% of the total number of nodes, and there should be fewer than 50 clusters

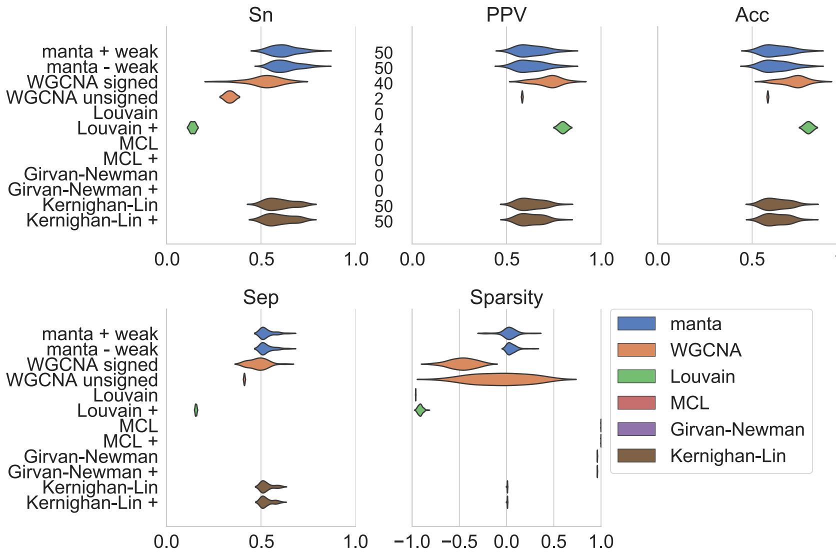


Figure S14: **Performance of clustering algorithms on networks with the range of edge weights shifted to 0 and 1.** Sensitivity (Sn), positive predictive values (PPV), accuracy (Acc) and separation (Sep) were calculated as described by [2]. Sparsity of the assignment is a function of the edge weights of intra-cluster versus inter-cluster edges (Equation 4). Clustering performance was estimated on 50 independently generated datasets. Matrices of taxon abundances were generated from a synthetic random interaction matrix. Clustering was carried out on Spearman correlation networks inferred from these matrices, with the range of correlations shifted to 0 and 1. As WGCNA constructed its own networks, shifting the edge weights was not possible for this tool and its performance therefore corresponds to performance on the normalized data. The numbers next to the sensitivity results indicate how many clustering assignments met the following criteria for a particular algorithm: no cluster should exceed 80% of the total number of nodes, and there should be fewer than 50 clusters.
